# Genetic basis of variation in thermal developmental plasticity for *Drosophila melanogaster* body pigmentation

**DOI:** 10.1101/2023.09.03.556093

**Authors:** E Lafuente, D Duneau, P Beldade

**Affiliations:** Instituto Gulbenkian de Ciência, Oeiras, Portugal; UMR5174, Laboratoire Évolution & Diversité Biologique, Université Paul Sabatier, Toulouse, France; Institute of Ecology & Evolution, University of Edinburgh, Edinburgh, UK; cE3c (Center for Ecology, Evolution and Environmental Changes) & CHANGE (Global Change and Sustainability Institute), FCUL, Lisboa Portugal

**Keywords:** developmental plasticity, thermal melanism, insect coloration, GWAS, DGRP, seasonal variation

## Abstract

Seasonal differences in insect pigmentation are attributed to the influence of ambient temperature on pigmentation development. This thermal plasticity is adaptive and heritable, thereby capable of evolving. However, the specific genes contributing to the variation in plasticity that can drive its evolution remain largely unknown. To address this, we analyzed pigmentation and pigmentation plasticity in *Drosophila melanogaster*. We measured two components of pigmentation in the thorax and abdomen: overall darkness and the proportion of length covered by darker pattern elements (a trident in the thorax and bands in the abdomen) in females from two developmental temperatures (17°C or 28°C) and 191 genotypes. Using a GWAS approach to identify the genetic basis of variation in pigmentation and its response to temperature, we identified numerous dispersed QTLs, including some mapping to melanogenesis genes (*yellow*, *ebony*, and *tan*). Remarkably, we observed limited overlap between QTLs for variation within specific temperatures and those influencing thermal plasticity, as well as minimal overlap between plasticity QTLs across pigmentation components and across body parts. For most traits, consistent with selection favoring the retention of plasticity, we found that lower plasticity alleles were often at lower frequencies. The functional analysis of selected candidate QTLs and pigmentation genes largely confirmed their contributions to variation in pigmentation and/or pigmentation plasticity. Overall, our study reveals the existence and underlying basis of extensive and trait-specific genetic variation for pigmentation and pigmentation plasticity, offering a rich reservoir of raw material for natural selection to shape the independent evolution of these traits.

## INTRODUCTION

The environment plays an important role not only in influencing changes in phenotype frequencies across generations, but also in shaping the expression of phenotypes within a single generation. Developmental plasticity, in particular, refers to the phenomenon where different phenotypes are produced based on the environmental conditions experienced during development (Beldade et al. 2011; Yoon et al. 2023). This plasticity becomes advantageous when the environmentally-induced phenotypes are better suited to the prevailing conditions compared to a fixed phenotype (Ghalambor et al. 2007; Nettle and Bateson 2015). As a result, developmental plasticity serves as an adaptive mechanism for organisms to effectively cope with environmental heterogeneity. A striking example of this is observed in cases of seasonal plasticity (Brakefield 1996; Brakefield and Zwaan 2011; Simpson et al. 2011), where alternative environmentally-induced phenotypes are tailored to distinct seasonal environments and enable season-specific strategies for survival and/or reproduction. Despite its ecological significance, we know very little about the genetic basis of variation in plasticity.

Insects, being ectotherms with relatively short life cycles, offer numerous examples of seasonal plasticity, as consecutive generations can experience the, often quite distinct, conditions of alternating seasons. Ambient temperature, a key cue associated with yearly seasons, undergoes significant fluctuations (Colinet et al. 2015), and thermal plasticity in insects has been extensively documented. Notably, various insect species exhibit remarkable thermal plasticity in body pigmentation (e.g. Gibert et al. 1996; Stoehr and Goux 2008; Sibilia et al. 2018; Lafuente et al. 2021), a trait crucial for interactions with both biotic factors (e.g., predator avoidance and mate choice) and abiotic factors (e.g., thermoregulation, UV protection, and desiccation resistance). Generally, insects display darker body pigmentation at lower temperatures, supposedly as an adaptive response to fulfill thermoregulatory needs (Clusella Trullas et al. 2007). While geographical clines in body pigmentation are attributed to adaptation to local environmental conditions, including temperature, seasonal variation in body pigmentation is typically attributed to developmental plasticity, often influenced by ambient temperature. For example, studies in *Drosophila* flies have documented that lower developmental temperatures result in adults with darker bodies (e.g. Gibert et al. 1996; Saleh Ziabari and Shingleton 2017; Lafuente et al. 2021). Additionally, melanization in *Drosophila* has been shown to be associated with thermoregulation and UV resistance (Matute and Harris 2013; Freoa et al. 2023), and correlated with various other traits, including behavior and immunity (Wittkopp and Beldade 2009; McKinnon and Pierotti 2010).

Studies in *Drosophila melanogaster* have played a pivotal role in advancing our understanding of the developmental, population, and evolutionary genetics of pigmentation (e.g. True 2003; Wittkopp et al. 2003; Gibert et al. 2007; Pool and Aquadro 2007; Massey and Wittkopp 2016; Gibert and Peronnet 2021). These investigations have also provided insights into the response of pigmentation development to temperature, including the characterization of thermal reaction norms (David et al. 1990; Gibert et al. 2000; Gibert et al. 2009) and an understanding of the molecular mechanisms underlying pigmentation plasticity (Gibert et al. 2007; Gibert et al. 2016; de Castro et al. 2018). However, despite our knowledge of genes involved in pigmentation development (True 2003; Wittkopp et al. 2003) and genes regulating thermal plasticity in pigmentation (Gibert et al. 2007; Gibert et al. 2016; Gibert et al. 2017; de Castro et al. 2018), we know very little about which genes harbor natural allelic variation contributing to variation in pigmentation plasticity. These genes represent potential targets for selection to drive the evolution of pigmentation plasticity in natural populations. The extent of overlap between genes involved in pigmentation development and those contributing to variation in plasticity remains unclear (Lafuente and Beldade 2019). Additionally, it is also unclear the extent of overlap between genes contributing to (inter-genotype) variation in plasticity and those contributing to (inter-individual) variation in pigmentation within specific temperature conditions, as well as whether the same genes contribute to variation in different plastic traits.

In this study, we investigated the genetic basis of thermal plasticity in body pigmentation, specifically focusing on the dorsal surface of the thorax and abdomen of *Drosophila melanogaster* flies. These body parts display distinct darker pattern elements, such as a trident on the thorax and bands on each abdominal segment, against a lighter background color. Pigmentation plasticity in these flies is characterized by the expansion of the darker bands in abdominal segments, particularly at the tip of the abdomen, resulting in an overall darker body appearance (e.g. Gibert et al. 2000; Gibert et al. 2009). To examine the genetic basis of variation in pigmentation and thermal plasticity therein, we used a method for quantitative analysis of *Drosophila* pigmentation (Lafuente et al. 2021). We evaluated the overall darkness and the proportion of each body part covered by darker pattern elements in adult female flies that developed at either 17°C or 28°C. Pigmentation and pigmentation plasticity were measured across 191 genotypes from the *Drosophila* Genetic Reference Panel (DGRP), a collection of fully-sequenced isogenic lines derived from a single natural population (Mackay et al. 2012; Huang et al. 2014). Our analysis revealed significant genetic and temperature effects on body pigmentation and enabled us to quantify genetic correlations among pigmentation components, body parts, and temperatures. Subsequently, we conducted a Genome-Wide Association Study (GWAS) on the pigmentation phenotypes (for flies developed at either temperature) and on their thermal plasticity (measured by the slope of thermal reaction norms). We identified multiple loci associated with variation in those traits, and we further characterized a subset of the corresponding candidate Quantitative Trait Loci (QTLs), as well as candidate genes known to be involved in development and variation in *Drosophila* pigmentation. Our results revealed substantial genetic variation but limited overlap in underlying loci between traits and between environmental contexts. This highlights the potential for independent evolution across body parts and pigmentation components, and between pigmentation and thermal plasticity therein.

## RESULTS

We investigated inter-genotype variation in pigmentation and pigmentation plasticity in *D. melanogaster* by analyzing overall darkness (Odk) and proportion of transect length covered by the darker pattern element (Pat) of fly abdomens (aOdk and aPat) and thoraxes (tOdk and tPat) (Figure 1). We phenotyped adult females from 191 isogenic lines developed at either 17°C or 28°C, and documented significant variation between genotypes and between temperatures for the four target traits (Figure 2). Additionally, we documented significant genotype-by-temperature interactions, corresponding to differences between genotypes in the magnitude and/or direction of temperature effects. Using these data and a GWAS approach, we identified loci contributing to inter-genotype variation in the slope of thermal reaction norms (RNs), as well as in body pigmentation at each of the two target temperatures (Figure 3, S2). Finally, using different fly lines and approaches, we validated the role of selected GWAS hits (Figure 4) and of known candidate pigmentation genes (Figure 5) in variation for body pigmentation and/or for pigmentation plasticity.

**Figure 1.**
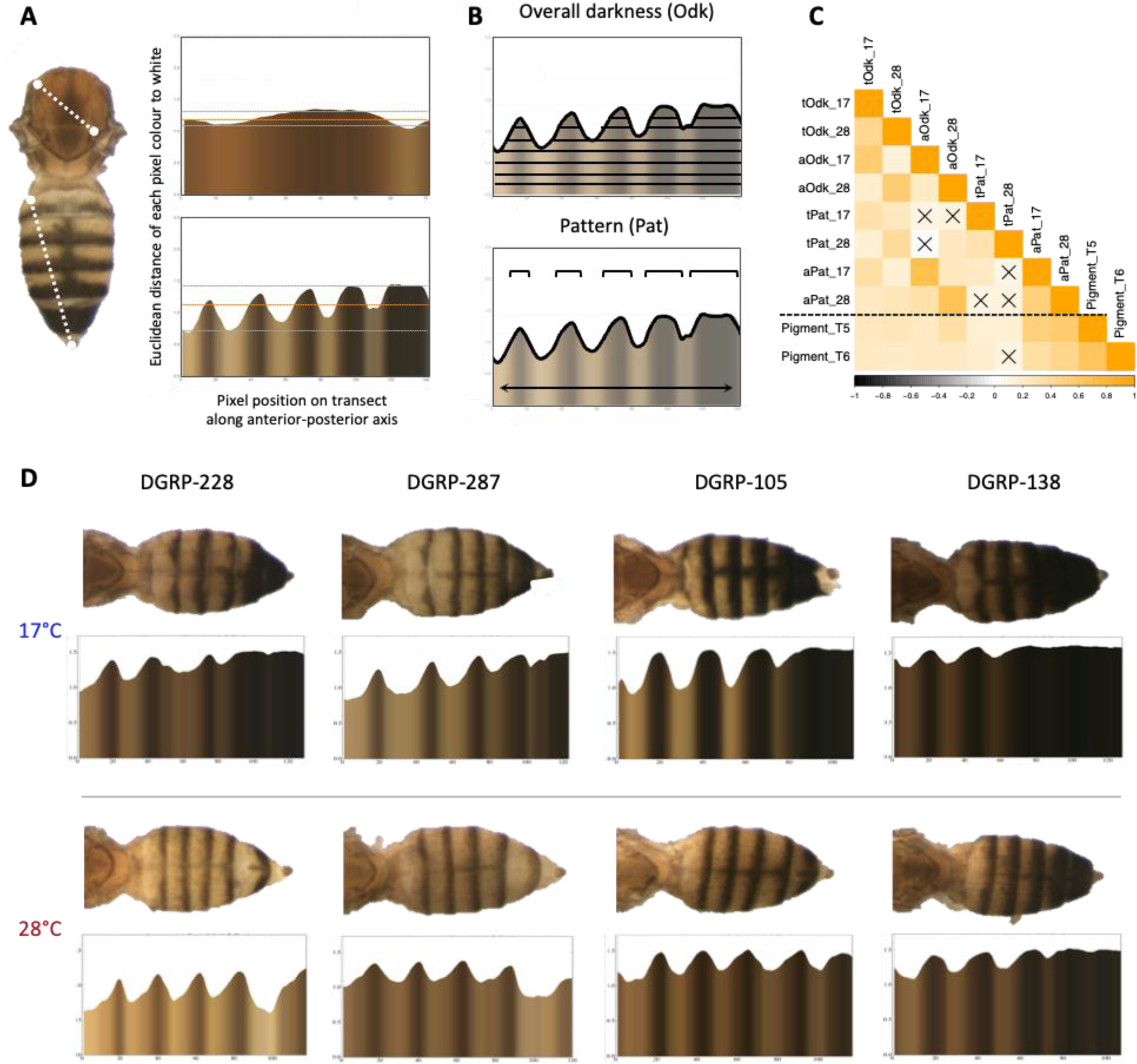
Pigmentation traits. **A)** Dorsal view of thorax and abdomen of mounted adult female fly, illustrating the transects drawn between body landmarks along which we extracted each pixel’s RGB values. For each body part, the plot corresponds to the distance between each pixel along the transect and the white vector (RGB 1,1,1). Details about transect position, processing of RGB data, and horizontal lines in plots can be found in the Materials & Methods section. **B)** Overall darkness (Odk) was calculated as the sum of the normalized darkness values for each pixel divided by the number of pixels in the transect. Pattern (Pat) was calculated as the proportion of pixels corresponding to the pattern element (thoracic trident or darker abdominal bands) relative to the transect length. **C)** Spearman correlations between pigmentation traits quantified in this study for the various DGRP genotypes (above dashed line), as well as between ours and abdominal pigmentation traits quantified in another study using the DGRP (Dembeck et al. 2015). Color gradient reflects Spearman correlation coefficient (positive in orange, negative in gray). Non-significant correlations (Holm-Bonferroni adjusted p-value > 0.05) are indicated with an ‘X’. **D)** Illustration of differences in abdominal pigmentation between genotypes (four DGRP lines) and developmental temperatures (17°C and 28°C), showing the fly image (dorsal view) on top and the corresponding plot (cf panel A) on the bottom.

**Figure 2.**
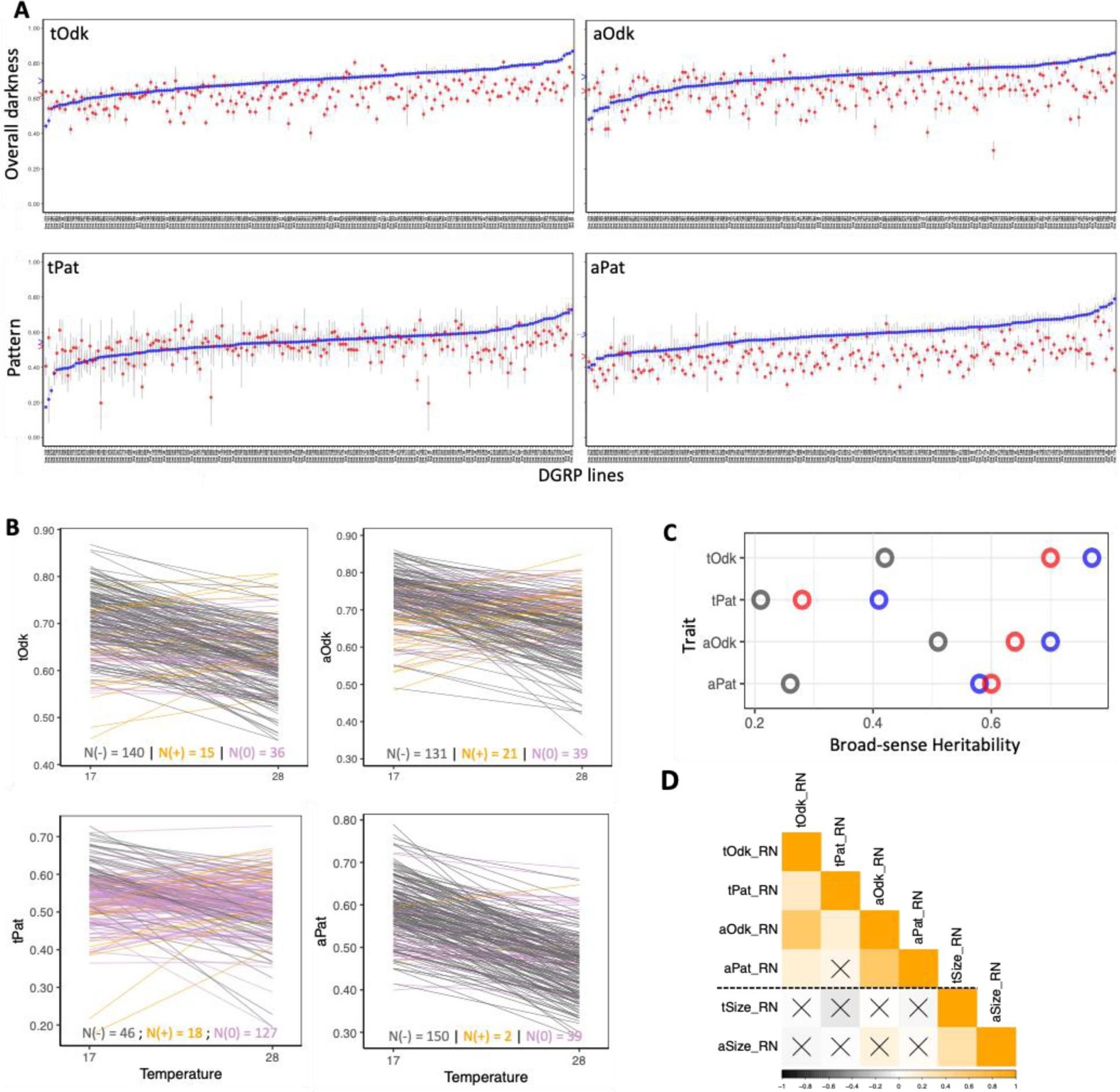
Genetic variation for pigmentation and thermal plasticity in pigmentation. **A)** Means and 95% confidence intervals of the pigmentation traits (tOdk, aOdk, tPat, aPat) for each of the target DGRP genotypes (X axis) reared at 17°C (blue symbol) and 28°C (red symbol). Genotypes are ordered by the mean phenotype at 17°C. Arrow heads next to Y axes represent the mean trait values for each of the temperatures. **B)** Reaction norms for target pigmentation traits (Y axis) as a function of developmental temperatures (X axis) corresponding to the regression fir for each genotype. Line color indicates sign of slope: positive slopes in orange, negative slopes in gray, and slopes not significantly different from zero (p-value>0.05) in purple. The number of genotypes in each of these classes is shown at the bottom of each plot. **C)** Values for broad-sense heritability (H^2^) of the four target traits (tOdk, aOdk, tPat, aPat) measured in flies from 17°C (blue) or 28°C (red), as well as for the slope of the reaction norms (gray). **D)** Spearman correlations for the slopes of thermal reaction norms across genotypes. Cells above the dashed line correspond to traits quantified in this study, while cells below the dashed line correspond to traits from another study that quantified slope of thermal reaction norms (Lafuente et al. 2018). Color gradient reflects Spearman correlation coefficient (positive in orange, negative in gray). Non-significant correlations (Holm-Bonferroni adjusted p-value > 0.05) are indicated with an ‘X’.

**Figure 3.**
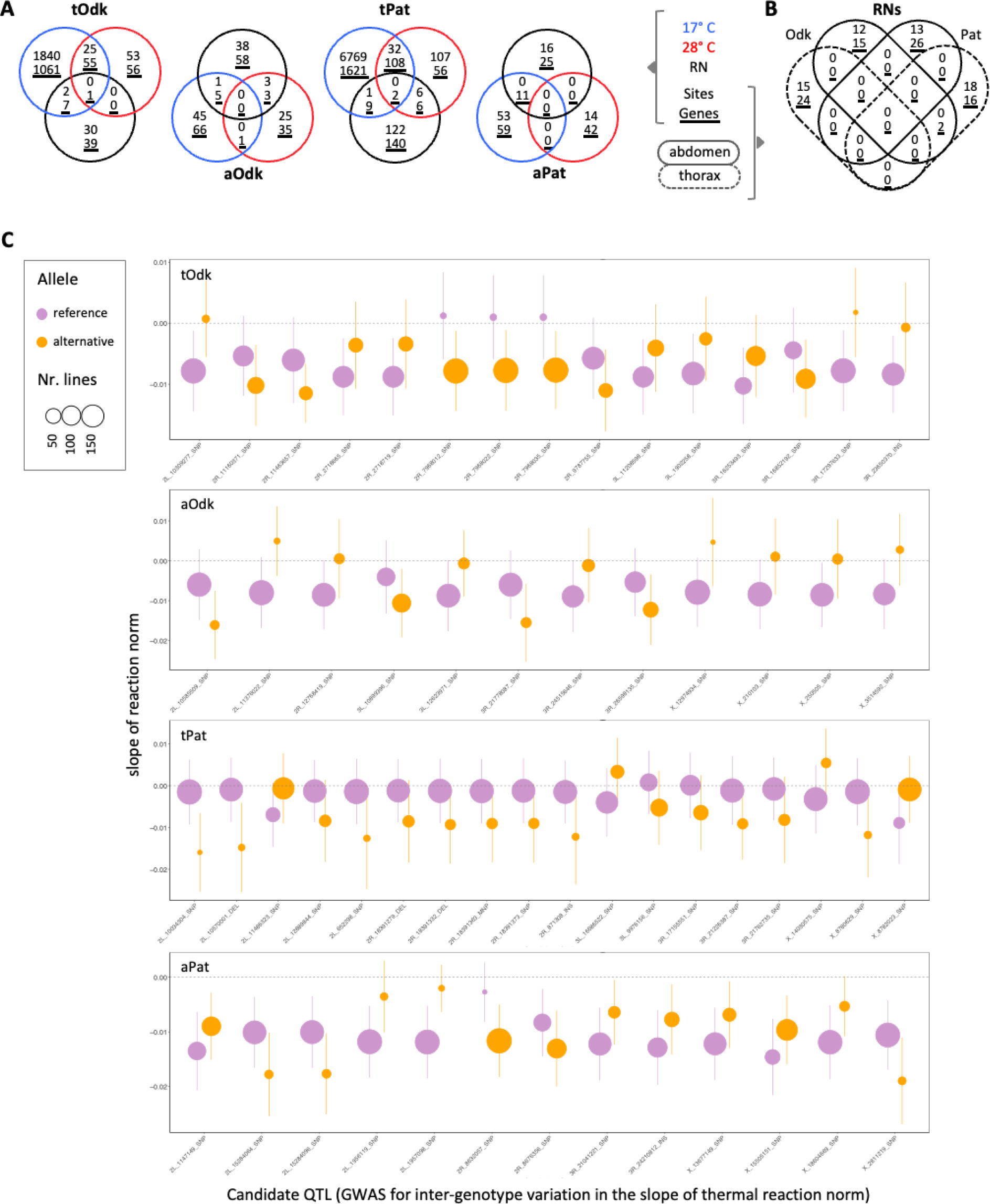
GWAS-generated candidate QTLs for variation in thermal plasticity in body pigmentation. **A)** Overlap in candidate QTLs for inter-genotype variation for body pigmentation in flies developed at 17°C (blue) or at 28°C (red), as well as for thermal plasticity in body pigmentation, considering both the raw value and absolute value of the slopes of thermal reaction norms (black). Venn diagrams show overlap in the identity of the polymorphic sites (SNPs/InDels) with a p-value<10e-5 (number on top), and the genes putatively associated to those sequence variants (underlined number below). **B)** Overlap in candidate QTLs for inter-genotype variation for plasticity in body pigmentation across traits (Odk to the left, Pat to the right) and across body parts (solid line for abdomen, dashed line for thorax). For each class, candidate QTLs correspond to SNPs/InDels with a p-value<10e-5 from the GWAS considering the raw value and the absolute value of the slopes of thermal reaction norms. Number of candidate polymorphisms is given on top, while number of genes these polymorphisms are putatively associated is given below and is underlined. **C)** Absolute frequency (size of circles) and effects (Y axis) for the alternative alleles (reference allele in purple, alternative allele in orange) for each of the significant QTLs associated with variation in the slope (raw values) of thermal reaction norms. The horizontal dashed line marks slope=0, corresponding to no plasticity.

**Figure 4.**
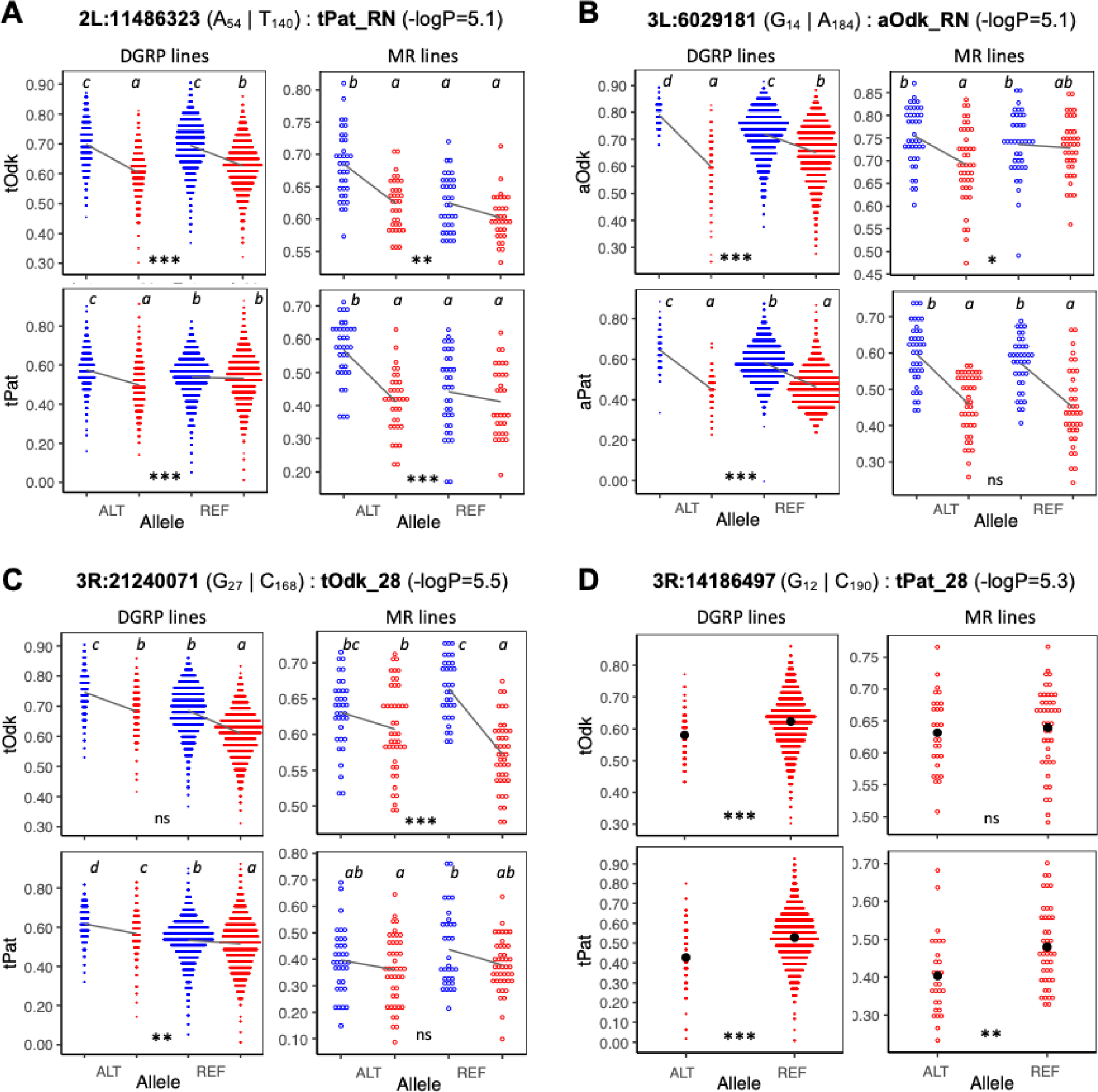
Mendelian Randomization validation of selected candidate SNPs from the GWAS analyses. For each candidate SNP, figures display phenotype of individual flies (one dot per individual fly) from either the DGRP lines (i.e. the data from the GWAS, left) or the Mendelian randomization lines (right). Flies are separated by rearing temperature (blue for 17°C, red for 28°C) and by allele at candidate SNP (REF for reference allele, ALT for the alternative allele in relation to reference genome). Black lines correspond to thermal reaction norms (linear regression). Inside each plot, italic letters on top represent pairwise statistical differences (significant if different letters; cf. Tukey’s honest test), while characters at the bottom represent significance of genotype-by-temperature interactions, corresponding to slope differences between reaction norms, or between trait values in the case of panel D: ns for p>0.05, * for p<0.05, ** for p<0.01, *** for p<0.001. Notation above plots provides information about each candidate SNP, including SNP identity (with chromosome and position location), alternative alleles (letters within brackets) and their absolute frequency (subscript), the trait the SNP was a GWAS hit for (after colon) and corresponding p-value (in brackets). **A)** SNP 2L:11486323, mapped to gene sala (missense variant) and to gene CG43355 (3’ UTR variant), was a significant hit in the GWAS for the raw value of the slope of reaction norms for tPat. **B)** SNP 3L:6029181, mapped downstream of genes PVRAP and CG10477, was a significant hit in the GWAS for absolute value of the slope of reaction norms for aOdk. **C)** SNP 3R:21240071, mapped upstream of genes ebony and CG5892, as a significant hit in the GWAS for tOdk at 28°C, but also for tOdk at 17°C (logP=5.4) and for tPat at 17°C (logP=10.0). **D)** SNP in position 3R:14186497, mapped to an intron of Dop1R1, was a significant hit in the GWAS tPat at 28°C.

**Figure 5.**
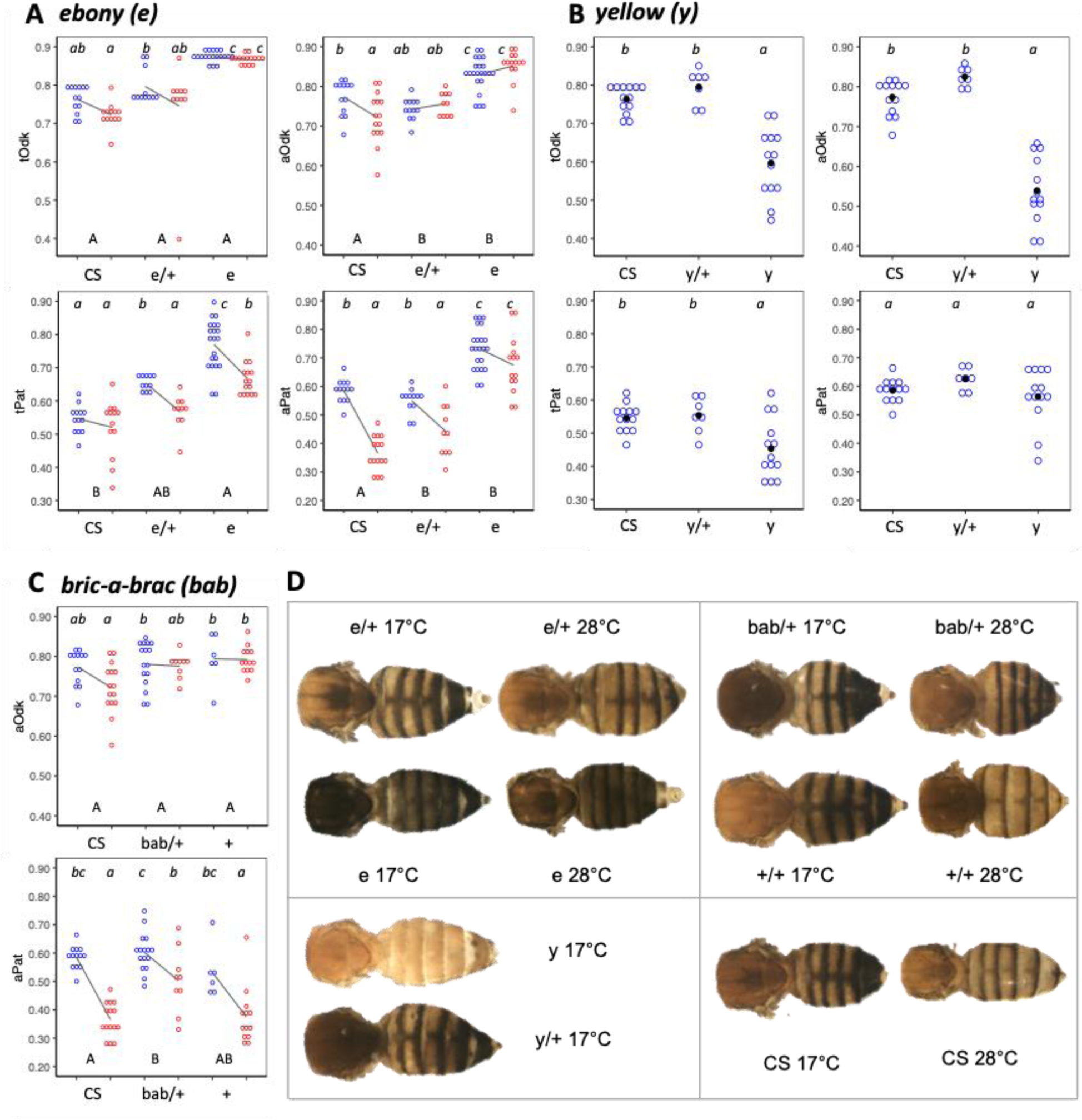
Role of candidate genes involved in pigmentation development. **A-C)** Each dot represents one individual female fly, with flies separated by genotype (Canton-S (CS), mutant homozygotes (e, y) or heterozygotes (e/+, y/+, bab/+)) and rearing temperature (blue for 17C and red for 28C). Back lines correspond to thermal reaction norms. In each plot, italic letters above represent pairwise statistical differences (significant if different letters; cf. Tukey’s honest test), while capital letters on the bottom represent differences between reaction norms (significant genotype-by-temperature interactions if different letters; cf. lm(Traits∼Genotype*Temperature), alpha=0.05). Three target genes implicated in pigmentation development: ebony (**A**), yellow (**B**; for which we only phenotype individuals from 17°C due to low survival at 28°C), and bric-a-brac1 (**C**; for which we only phenotyped abdomens). **D)** Representative images of dorsal view of female thoraces and abdomens from different genotypes and developmental temperatures.

### Genetic and environmental variation for body pigmentation

We phenotyped 5006 female flies (Table S1) and found significant differences between genotypes (G) and between rearing temperatures (T) for our four target traits (lm, with p<1e-14 in all cases; see Table S2): tOdk (G: F_199,4356_=48.6, T: F_1,4356_=3300.6), aOdk (G: F_199,4521_=30.9, T: F_1,4521_=1980.1), tPat (G: F_199,4356_=8.3, T: F_1,4356_=61.6), aPat (G: F_199,4521_=32.0, T: F_1,4521_=4573.8) (Figure 2A). Broad-sense heritability (H^2^) estimates, calculated based on levels of within and between line variance (see Materials and Methods), ranged from 0.28 (for tPat at 28°C) to 0.77 (tOdk at 17°C), and were generally higher for Odk than for Pat and higher for 17°C than for 28°C (Figure 2C). There were significant positive correlations between most pairs of traits across the two temperatures, with all exceptions involving trait tPat (Figure 1C). Additionally, we found significant positive correlations between our body part-wide measurements of pigmentation in flies reared at 17°C and 28°C and previously characterized segment-specific pigmentation (abdominal tergites T5 and T6) in flies reared at 25°C (Dembeck et al. 2015), with only one exception (Figure 2A).

We confirmed typical patterns of temperature effects on insect pigmentation, with flies from lower developmental temperature generally being darker than those from higher temperature (Figure 1D), reflected in all four traits we quantified (Figure 2A, Figure S1A). This is consistent with global patterns described for *Drosophila* flies (David et al. 1990; Gibert et al. 2000; Lafuente et al. 2021), and believed to be associated with thermoregulation needs (Clusella Trullas et al. 2007; Freoa et al. 2023).

### Inter-genotype variation in magnitude and direction of pigmentation plasticity

We found significant genotype-by-temperature (GxT) interactions for all traits (aOdk: F_190,4521_=17.6, aPat: F_190, 4521_=9.0, tOdk: F_190,4356_=17.6, tPat: F_190,4356_=4.5; p<1e-14 in all cases; Table S2), corresponding to non-parallel thermal reaction norms across genotypes (Figure 2B). Estimates of broad-sense heritability for GxT effects varied between 0.21 (for tPat) and 0.51 (for aOdk). These were lower than estimates of heritability for within temperature variation, and were again higher for Odk than for Pat (Figure 2C). For most genotypes, flies reared at 17°C were darker (higher values of Odk and of Pat) than flies reared at 28°C. This is the general trend for thermal plasticity in body pigmentation across Drosophila species (Lafuente et al. 2021).

However, this was not the case for all genotypes we analyzed (Figure 2B). Some genotypes had reaction norm (RN) slopes that were not significantly different from zero, corresponding to no difference between flies from 17*°*C versus 28*°*C (i.e. no thermal plasticity): around 20% for tOdk, aOdk, and aPat RNs, and 66% for tPat RNs. There were also genotypes with positive RN slopes, corresponding to darker bodies for flies from higher rearing temperature; ranging from 1% for aPat to 11% for aOdk. Of these, only nine genotypes had “reversed plasticity” (i.e. positive instead of the more common negative reaction norm slopes) for more than one of the target traits. Thermal plasticity in the opposite direction of the general trend in insects had been documented for Drosophila body size in the same DGRP lines, where not all genotypes produced larger bodies when developed at lower temperature (Lafuente et al. 2018).

We found significant positive correlations between the slopes of thermal reaction norms across pigmentation traits, except between tPat and aPat, but no significant correlations with the slopes of thermal reaction norms for thorax and abdomen size measured in another study (Lafuente et al. 2018) (Figure 2D). This suggests a different basis for thermal plasticity in different types of traits. Consistent with the trait-specific responses is the observation that reaction norms often differ between traits (e.g. Gibert et al. 2009; Oostra et al. 2011; Mateus et al. 2014).

### Genetic basis of inter-genotype variation in pigmentation and pigmentation plasticity

For each of our four target pigmentation traits (tOdk, tPat, aOdk, aPat), we performed four GWAS using distinct measurements. We analyzed pigmentation in flies from each of the two rearing temperatures (variation between individuals reared at 17*°*C and variation between individuals reared at 28*°*C), as well as two measurements of thermal plasticity in pigmentation, the raw and absolute values of the slopes of thermal reaction norms (Figure 4 and Figure S2). The raw value of slopes reflects both the direction and magnitude of the plastic response, while the absolute value reflects thermal sensitivity, describing only the magnitude of the response to temperature. This approach allowed us to identify QTLs for pigmentation at 17*°*C and 28*°*C and for thermal plasticity in pigmentation. The results of the GWAS analyses are detailed in Figure S2 (Manhattan Plots) and Table S3 (list and details of significant candidate QTLs), including which genes they are putatively associated to and gene regions they fall within (*cf.* Genome assembly release 6; dos Santos et al. 2015). Rather than Manhattan Plots (Figure S2) with a small number of clear peaks composed of neighboring high significance SNPs, consistent with a small number of QTLs of stronger effect, we found numerous and dispersed QTLs. For thorax pigmentation in flies from 17°C, there was an excess of significant hits concentrated on chromosome 3R (1841 out of a total 1920 for tOdk, and 6886 out of 6909 for tPat; Table S3, Figure S2), most of which are likely not real hits. The same was not observed for the GWAS results for variation in abdominal pigmentation or for variation in pigmentation plasticity in either body part.

The annotation of the Drosophila genome allowed us to establish a correspondence between a DNA sequence polymorphism and one or several genes it might potentially affect (Table S3). Our GWAS hits included SNPs potentially mapping to genes with well-documented roles on pigmentation biosynthesis and/or pigmentation plasticity, including *ebony* (e), *yellow* (y), and *tan* (Massey and Wittkopp 2016; Gibert et al. 2016, 2017), but did not include *bric-a-brac* (*bab1*), which has been implicated in regulation of pigmentation development (Kopp et al. 2000) and plasticity (de Castro et al. 2018) and was a candidate QTL for variation in pigmentation of abdominal segments T5 and T6 (Dembeck et al. 2015). In common with the QTLs identified in this study, which investigated the genetic basis of pigmentation variation in DGRP flies reared at 25°C, we found 14 candidate sequence polymorphisms and 34 candidate genes. However, overall, we found that QTLs for pigmentation variation were largely temperature specific. For example, for aOdk and aPat, traits for which we did not observe the potential artefactual excess of QTLs in chromosome 3R (Figure S2), there was no single QTL in common between variation at 17*°*C and variation at 28*°*C (Figure 3A).

To address whether the genes contributing to inter-genotype variation in thermal plasticity are the same genes that account for variation within each temperature, we calculated the extent of overlap between QTLs for the slope of thermal reaction norms and the QTLs for pigmentation at 17°C of 28°C (Figure 3A). To address whether the genes contributing to variation in plasticity are the same across pigmentation components and across body parts, we calculated the overlap between QTLs for the slope of reaction norms across our four pigmentation traits: tOdk, aOdk, tPat, and aPat (Figure 3B). Globally, we found that few candidate QTLs were common between different GWAS analyses; i.e. most QTLs identified seemed “private” rather than shared. There was very little overlap between QTLs contributing to variation in plasticity (identified from GWAS on slopes of reaction norms) and QTLs contributing to within-environment variation (identified from GWAS on pigmentation values for either 17*°*C or 28*°*C) (Figure 3A). Curiously, the single exception of a SNP associated with both variation in tPat at 17*°*C and variation in tPat plasticity was SNP *X:356472*, mapped upstream of the melanogenesis gene *yellow*. Additionally, our analysis revealed that plasticity QTLs were trait-specific (i.e. little overlap between Odk and Pat) and body-part-specific (i.e. little overlap between thorax and abdomen) (Figures 3B). There were only two cases of the same gene (but not the same SNP) contributing to variation in plasticity for tPat and aOdk (genes *Trim9* and *CCAP-R*). These results, of trait, body part, and environment specific QTLs, matches what had been described for inter-genotype variation in body size and thermal plasticity therein (Lafuente et al. 2018).

To assess whether alleles associated with increased plasticity are potentially favored by natural selection, we plotted allelic effect and frequency for the QTLs for variation in slope of thermal reaction norms (Figure 3C). For tOdk, aOdk, and aPat, alleles for reduced plasticity (i.e. closer to the horizontal line of no plasticity in Figure 3C) were often, albeit not always, at lower frequency among DGRP genotypes (i.e. they were the minor allele, represented by the smaller circles in Figure 3C). Additionally, all cases with reversed plasticity (i.e. positive slopes of reaction norms, which are generally negative) were also minor alleles. This contrasts what was described for thermal plasticity in body size in the DGRP (Lafuente et al. 2018). The higher frequency for alleles corresponding to higher plasticity is consistent with pigmentation plasticity being favored by selection in the population the DGRPs were derived from, and does not support the existence of potential high costs for pigmentation plasticity that could have led to its loss in the DGRP lines. For tPat, the results were quite different. For all candidate QTLs for variation in tPat plasticity, which had lowest value for broad-sense heritability (Figure 2C), the allele corresponding to lower plasticity (slope closer to zero) were at high frequency among DGRP genotypes (i.e. they were the major allele) (Figure 3C).

### Analysis of selected candidate SNPs and candidate genes

We selected four SNPs from our GWAS hits (corresponding to QTLs for different traits), and three genes known to be involved in body pigmentation development (*e*, *y*, *bab1*) for further targeted analysis. We used a SNP-centered Mendelian Randomization (MR; cf. Lafuente et al. 2018) method to validate candidate QTLs from the GWAS (Figure 4), and a gene-centered method using available null mutants to study effects of the pigmentation genes (Figure 5).

The MR approach tests whether individuals with one versus the other allele at the candidate SNP differ in the quantitative trait the QTL was identified for when in a novel, mixed genetic background. For each of the candidate SNPs, the MR approach involved randomizing the genetic background between 10 same-allele genotypes and then comparing the quantitative trait between flies carrying the reference versus the alternative allele (see Material and Methods). We used the new lines to test for phenotypic differences between alternative alleles (G effect) and between alternative temperatures (T effect), and for differences in thermal plasticity (interaction between the G and T effects). Significant T effects reveal thermal plasticity, significant G effects reveal inter-genotype differences in body pigmentation, and significant GxT effects reveal inter-genotype differences in pigmentation plasticity. For all SNPs tested using this approach, we confirmed the association between allelic variants and the trait for which the SNP was identified as QTL: SNP *2L:11486323* (notation: chromosome:position *cf. Drosophila* genome v6) for plasticity in tPat (Figure 4A), SNP *3L:6029181* for plasticity in aOdk (Figure 4B), SNP *3R:21240071* for tOdk at 28°C (Figure 4C), and SNP *3R:14186497* for tPat at 28°C (Figure 4D). Note the association between SNP *3R:21240071* and tOdk and tPat at 17°C identified for the DGRP lines (Table S3), which was not validated in the MR lines (Figure 4C), was part of the region with excess QTLs in chromosome 3R (Table S3), which we believe to include artifactual associations. In most cases (Figure 4B-D), we confirmed trait-specific allelic effects. For example, SNP *3L:6029181* (Figure 4B), which was identified as a QTL for aOdk but not aPat plasticity in the DGRP lines, showed an association with plasticity variation for aOdk but not for aPat in the MR lines. On the other hand, SNP *2L:11486323*, identified as a QTL for tPat but not tOdk plasticity, was associated with differences in plasticity for both tPat and tOdk in the MR lines (Figure 4A).

Taking advantage of the genetic resources available for *D. melanogaster*, we tested the contribution of candidate pigmentation genes to pigmentation plasticity by comparing phenotypes between individuals with normal versus abolished protein function. We analyzed three candidate genes known to be involved in pigmentation development and to contribute to pigmentation variation in *Drosophila*; two genes encoding enzymes involved in melanogenesis (*ebony* and *yellow*; Figure 5A and Figure 5B) and a gene encoding the transcription factor *bric-a-brac1* (Figure 5C) that regulates the expression of melanogenesis genes and is involved in pigmentation development, evolution, and plasticity (e.g. Kopp et al. 2000; Rogers et al. 2013; Massey and Wittkopp 2016; de Castro et al. 2018). Our GWAS results identified candidate QTLs (Table S3) putatively mapping to *yellow* (SNP *X:356472* for tPat at 17°C and tPat plasticity) and to *ebony* (SNP *3R:21240071* for tOdk at 28°C and 17°C and tPat at 17°C, as well as 27 additional sequence polymorphisms accounting for variation in tOdk and/or tPat at 17°C; Table S3). Previous GWAS results had implicated *bab1* in variation in pigmentation of abdominal segments T5 and T6 in flies reared at 25*°*C (Dembeck et al. 2015). Consistently with what is well known about the function of these genes, loss of function of *ebony* resulted in darker flies and loss of function of *yellow* in lighter ones (Figure 5D), corresponding to high (Figure 5A) and low Odk values (Figure 5B), respectively. Except for tPat, we did not find significant differences between 17*°*C and 28*°*C *ebony* mutant flies, which corresponds to there being no thermal plasticity. Due to high mortality, we did not have enough adult mutant *yellow* flies from 28*°*C and could not assess thermal plasticity. Interestingly, even though *yellow* flies from 17*°*C had lighter abdomens (lower aOdk), they did not have thinner abdominal bands (no difference in aPat). For *bab1*, we phenotyped abdomens and found differences between flies carrying one versus none loss-of-function *bab1* allele for aPat at 28*°*C, but not for aOdk nor for plasticity in either trait.

## DISCUSSION

We found clear G, T, and GxT effects on body pigmentation among the wild-derived and isogenized *Drosophila melanogaster* lines from the DGRP (Figure 1D, Figure 2A, 2B). These effects correspond to differences between genotypes, between rearing temperatures, and between how genotypes respond to temperature (i.e. thermal plasticity), respectively. We estimated broad-sense heritability above 0.4 for most traits measured; in particular for pigmentation variation within temperature (Figure 2C). For the majority of genotypes we analyzed, lower developmental temperatures produced darker bodies (Figure 2B). Darker flies have been shown to have better light absorption (Freoa et al. 2023) and better protection against UV radiation (Bastide et al. 2014, but see Rajpurohit et al. 2019). However, some genotypes had no thermal plasticity (no difference in phenotype across temperatures) and a few had even inverted plasticity (with darker bodies at higher temperatures). While this type of “reversed plasticity” has been documented for pigmentation in other insects (Yin et al. 2015) and for other traits in *D. melanogaster* (Lafuente et al. 2018), it is unclear whether and under what conditions it would be favored by natural selection in wild populations. Darker bodies at higher temperatures defies finding of pigmentation evolution in warmer climates (Zeuss et al. 2014; Stelbrink et al. 2019) and, more generally, the hypothesis of thermal melanism proposed for ectotherms (Clusella Trullas et al. 2007), which emphasizes the association between body coloration and thermoregulation needs. However, pigmentation is involved in ecological functions other than thermo-regulation (Bastide et al. 2014; Gibert et al. 2018; Massey et al. 2019) and its expression is affected by environmental cues other than temperature (Shakhmantsir et al. 2014). As such, it is difficult to predict how environmental perturbation, which might disrupt association between environmental variables, will affect the evolution of pigmentation and pigmentation plasticity.

Using our DGRP pigmentation data at 17*°*C and 28*°*C and the slope of thermal reaction norms as quantitative traits in a GWAS, we identified sequence polymorphisms significantly associated with variation in those traits. We found a large number of candidate QTLs distributed across the fly genome (Figure S2, Table S3), which holds for all traits, even after we exclude the potentially artefactual QTLs for thoracic pigmentation in flies from 17*°*C concentrated on chromosome 3R (Figure S2). GWAS hits include known melanogenesis genes *yellow, ebony*, and *tan*, known to contribute to pigmentation development and evolution in *Drosophila* (Wittkopp et al. 2002; True et al. 2005; Massey and Wittkopp 2016) and to pigmentation plasticity (Gibert et al. 2016, 2017) in *D. melanogaster*. However, hits did not include other expected candidates such as the transcription factor *bric-a-brac*, involved in abdominal pigmentation plasticity (de Castro et al. 2018), and shown to contribute to pigmentation variation in abdominal tergites T5 and T6 in another GWAS study of DGRP flies (Dembeck et al. 2015). Our analysis of loss-of-function mutants for candidate genes *ebony*, *yellow,* and *bab1* provided quantitative data for our different pigmentation traits (Figure 5): while *ebony* affects both Pat and Odk, *yellow* affects mostly Odk, and *bab1* affects mostly Pat.

Despite significant positive correlations between our study traits (Figure 1C, 2D), the overlap in candidate QTLs underlying inter-genotype variation in each of them was minimal (Figure 3A). This result has important technical and conceptual implications. First, the limited overlap between QTLs for pigmentation across developmental temperatures (e.g. no single common QTL for variation in aOdk between 17*°*C and 28*°*C; Figure 3A) highlights the environmental-specificity of GWAS-derived QTLs and raises the concern that comparisons between studies are compromised. Second, the fact that QTLs for variation in pigmentation at 17*°*C or 28*°*C are largely not the same as QTLs for pigmentation plasticity (e.g. no common aPat QTL between 17*°*C or 28*°*C and plasticity; Figure 3A) sheds light into controversy around the existence of “plasticity genes” beyond genes responsible for variation in trait means (Via et al. 1995; Pigliucci 1998) and whether selection can target plasticity directly or whether plasticity is necessarily a by-product of selection for trait phenotype under different environments (Via 1993; Lafuente and Beldade 2019).

Similarly, despite significant positive correlations between our plasticity measurements (Figure 2C), we also found minimal overlap between plasticity QTLs across body parts (i.e. QTLs for plasticity in thorax pigmentation are largely not the same as QTLs for plasticity in abdominal pigmentation) and across pigmentation components (i.e. QTLs for plasticity in Odk are largely not the same as QTLs for plasticity in Pat) (Figure 3B). There was no single polymorphic site (and only two genes) in common between the QTLs for the slope of the different reaction norms analyzed. Moreover, and consistent with this, there was also very limited overlap (one single SNP, *3L:7461922*) in the QTLs for pigmentation plasticity identified here and the QTLs for size plasticity identified in a previous study (Lafuente et al. 2018). Limited overlap between genes contributing to variation in thermal plasticity for different traits argues against the existence of organism-wide plasticity QTLs that could drive the concerted evolution of plasticity for all plastic traits. While this might provide potential for much flexibility in the evolution of plasticity, it also has the potential to break the integration of suites of functionally related traits that change in concert in response to environmental input and make up what are often called “plasticity syndromes” (Brakefield and Zwaan 2011; van Bergen et al. 2017). Assessing the relevance of plasticity to fitness and to how organisms cope with environmental perturbation requires consideration of whole organism plasticity instead of focus on individual plastic traits (Forsman 2015).

Plasticity in pigmentation and in other traits is a heritable trait that can evolve (Scheiner 1993; Ørsted et al. 2018; Lafuente and Beldade 2019) and QTLs for plasticity provide the raw material and targets for natural selection to drive its evolution. We found numerous candidate QTLs for thermal plasticity in pigmentation in the DGRP lines. Additionally, we found that alleles associated with lower plasticity were often present at lower frequencies, consistent with retention of plasticity being favored by selection. However, it is unclear to what extent the alleles represented in DGRP lines reflect and/or could feed the evolution of plasticity in this and other natural populations. Moreover, a number of recent studies of multiple natural and semi-natural populations of *D. melanogaster* have revealed that seasonal variation in body pigmentation and other traits, which is generally attributed to thermal plasticity, can also be the result of fast genetic adaptation (e.g. Bergland et al. 2014; Behrman et al. 2018; Rudman et al. 2022). Partitioning the contribution of seasonal adaptation versus seasonal plasticity will undoubtedly be an active topic of analysis of data generated by various large-scale collaborative studies that are ongoing (Kapun et al. 2020; Kapun et al. 2021; Machado et al. 2021). Additionally, future research will also certainly continue to explore whether and how thermal plasticity and plasticity genes are affected by temperature perturbation resulting from climate change (Kingsolver and Buckley 2017; Sandoval-Castillo et al. 2020; Svensson et al. 2020), and, conversely, what their role might be in the response of natural populations to climate change (Merilä and Hendry 2014; Sgrò et al. 2016; Bonamour et al. 2019; Fox et al. 2019; Kelly 2019; Rodrigues and Beldade 2020).

## MATERIALS AND METHODS

### Fly stocks and rearing conditions

We analyzed adult female flies from different genotypes for the project’s different tasks. For the characterization of pigmentation variation, we used genotypes from the Drosophila Genetic Reference Panel (DGRP) (Mackay et al. 2012; Huang et al. 2014), which we obtained from the Bloomington Stock Center. We obtained data on flies reared at 17°C and 28°C from 191 DGRP genotypes, and data for only 17°C for three additional DGRP genotypes (for which survival was low at 28°C). For a subset of 75 DGRP lines at 17°C and 85 DGRP lines at 28°C, we generated two or three replicate sets. For the functional validations, we used: i) mixtures of selected DGRP genotypes, which we derived as described below (section “Functional analysis of candidate SNPs by Mendelian Randomization”), ii) loss-of-function mutants of selected pigmentation genes (*ebony*, *yellow*, and *bric-a-brac*), which we obtained from the Bloomington Stock Center (reference codes 1658, 169, and 37298, respectively), and iii) a standard control “wildtype” laboratory strain, Canton-S, which we obtained from CK Mirth lab. All fly stocks were maintained in incubators at 25°C, 12:12 light cycles, and 65% humidity, and fed *ad libitum* with molasses food (45 gr molasses, 75 gr sugar, 70 gr cornmeal, 20 gr Yeast extract, 10 gr Agar, 1100 ml water and 25 ml 10% Niapagin). For the experiments, we collected between 20 and 40 eggs from overnight egg lays of ∼20 females of each stock (in vials). Eggs were then transferred to either 17°C or 28°C to complete development. The total number of flies obtained per genotype and temperature varied due to line- and temperature-specific rearing success.

### Phenotyping body pigmentation

Processing and phenotyping flies was done as described in Lafuente et al. 2021. In short, 8-10 days after adult eclosion, live females were placed in 2-ml microcentrifuge tubes, submersed in liquid nitrogen (2-3 seconds) and immediately shaken to remove wings, legs, and bristles. Headless bodies of flies were mounted (dorsal side up) on Petri dishes containing a thin layer of 3% agarose, and covered with water to avoid specular light reflection during imaging. Images were collected with a LeicaDMLB2 stereoscope and a Nikon E400 camera under controlled conditions of illumination and white-balance adjustment, and were later processed with a set of custom-made interactive Mathematica notebooks (Wolfram Research, Inc., Mathematica, version 10.2, Champaign, IL, 2015) to extract pigmentation measurements. We used morphological landmarks to define one thoracic and one abdominal transect on each fly (Figure 1A). For each pixel position along the transect line we calculated the mean RGB (red, green, blue) values of the closest five pixels located on a perpendicular line centered on the transect. Where the membranous tissue between segments was visible, corresponding transect sections were removed. To calculate our pigmentation traits from the RGB values, we first calculated a normalized darkness value using the formula (Dmax-Dbk)/Dmax for each pixel, where Dmax is the largest possible Euclidean distance between two colors in the RGB color space (in this case, Dmax=√3), and Dbk is the distance of the pixel’s color coordinates to the color black (R=0, G=0, B=0). “Overall darkness” (Odk) was computed as the sum of the normalized darkness values for all pixels divided by the number of pixels in the transect. Second, we calculated the Euclidean distance of each pixel’s color coordinates to those of white (R=1, G=1, B=1) and then estimated the enveloping lines of the upper and lower extremes of these values (upper and lower gray lines in Figure 1A) by calculating the baselines of the original and negated values using the Statistics-sensitive Non-linear Iterative Peak-clipping (SNIP) algorithm (Ryan et al., 1988). Using the median line of this envelope (horizontal orange line in Figure 1A) we separated the transect pixels into two clusters: pixels above median line corresponding to the darker pattern element (trident in the thorax and bands in the abdomen) and pixels below median line corresponding to the lighter background. “Pattern” (Pat) was calculated as the proportion of pixels corresponding to the pattern element relative to the transect length. An illustration of what Odk and Pat correspond to can be found in Figure 1B.

### Statistical analyses of G and T effects on pigmentation

All statistical analyses were conducted in R v 3.6.2 (Team 2019) (R Development Core Team 2017) and Rstudio (RCore Team 2018; RStudio Team 2016). The *tidyverse* R package suite (Wickham et al. 2019) was used for preparing all analyses and plots. The packages *lme4* (Bates et al. 2014) and *lmerTest* (Kuznetsova et al. 2017) were used to analyze linear mixed-effects models, *VCA* (Schuetzenmeister and Dufey 2022) to calculate variance components, *ape* (Paradis and Schliep 2019), and *phytools* (Revell 2012) to estimate phylogenetic signals.

We checked assumptions of parametric tests by visualizing QQplots and residuals’ plots and by using Shapiro test (for normality) and Bartlett test (for homoscedasticity). For each body part and pigmentation trait, we used linear models to test for differences across DGRP genotypes (in R syntax: *lm(Trait∼Genotype)*) or for the interaction between genotype and temperature (*lm(Trait∼Genotype + Temperature + Genotype:Temperature)*). Reaction norms for each DGRP line were obtained using the regression model l*m(Trait∼Temperature),* from which we extracted two properties of the reaction norms for each body part: the absolute value of the slope (as a measurement of thermal sensitivity, that describes only the magnitude of the response to temperature) and the raw value of the slope (as a measurement that describes also the direction of that response).

Broad-sense heritability (H^2^) for each pigmentation component and at each temperature was estimated as *H*^2^=*σ*^2^ /(*σ*^2^ +*σ*^2^), where *σ*^2^ and *σ*^2^ are the among-line and within-line variance components, respectively. Heritability of plasticity for each pigmentation component was calculated cf. Scheider and Lyman (1989): *H*^2^=*σ*^2^ /*σ*^2^, where *σ*^2^ and *σ*^2^ are the variance associated with the *genotype by environment* interaction component and the total variance, respectively. For the functional analysis of within-environment SNPs and genes, we used the models *lm(Trait∼Allele)* and *lm(Trait∼Genotype)*, respectively. For the functional analysis of plasticity SNPs and genes, we used the models *lm(Trait∼Genotype + Temperature + Genotype*:Temperature)* and *lm(Trait∼Allele + Temperature + Allele:Temperature)*, respectively. In all cases where there were more than two categories, significant pairwise differences were identified by post hoc Tukey’s honest test.

Genetic distance matrix for the DGRPs was obtained from http://dgrp2.gnets.ncsu.edu/data.html and was used to perform a cluster hierarchical and to estimate the phylogenetic signal and statistical significance for each of our traits using Blomberg’s K (Blomberg et al. 2003) and Pagel’s λ (Pagel 1999) metrics.

### Genome-Wide Association Study

For each pigmentation trait (tOdk, aOdk, tPat, aPat), we performed four independent genome wide analyses (GWAS): two for thermal plasticity (raw and absolute values of the slopes of the reaction norms used as target quantitative trait), and two for within-temperature variation (pigmentation at 17°C and 28°C as quantitative traits). For each polymorphic site (SNPs and InDels) with a minimum minor allele frequency of ten (*i.e.* minimum of 10 DGRP lines with each allele), we tested the models differences between alleles taking account of whether lines were or not infected with *Wolbachia*, corresponding to the R syntax: *lm(Slope∼Allele+(1|Wolb/DGRP))* or *lm(Trait∼Allele+(1|Wolb/DGRP)),* for variation in plasticity or variation within-temperature, respectively. *Allele* was used as a fixed factor with two values (*i.e.* reference versus alternative allele) and *Wolb* as a random factor corresponding to the *Wolbachia* status of each DGRP line (*i.e.* present or absent). We did not find an effect of *Wolbachia* on any of our GWAS analyses. We also did not find an effect on *Slope* or *Trait* of any of the chromosomal inversions present in at least 8 DGRP lines (In_3R_K, In_3R_P, In_2L_t, In_2R_NS and In_3R_Mo), by testing the models lm(*Slope∼Inversion)* and lm(*Trait∼Inversion)*, where Inversion is a fixed factor with two values (with and without inversion). For each of the GWAS analyses, we annotated the SNPs with a p-value < 10e-5 using the FlyBase annotation (release 6; dos Santos et al. 2015).

### Functional analysis of candidate SNPs by Mendelian Randomization

The contribution of selected GWAS hits was assessed using a method that we call Mendelian Randomization (MR; see Lafuente et al. 2018). We selected SNPs with p-value < 10e-5 based on their position in Manhattan plots, putative effect (missense and regulatory variants prioritized over inter-genic variants), and associated genes (annotated and known function prioritized). Validations by MR were done by selecting for each candidate SNP, 10 DGRP lines with the minor allele and 10 with the major allele, that were not fixed for any other significant SNP. These lines were used to generate four populations, two fixed for the reference allele and two for the alternative allele. Each population was established by crossing 8 virgin females from each of 5 of the same-allele lines to 8 males of the other 5 lines. The reciprocal crosses were used to set two independent populations per allele. These populations were allowed to cross for eight generations to randomize genetic backgrounds. We confirmed by Sanger sequencing of target amplicons that those populations had our candidate allele fixed.

## Author contributions

Study conception: EL, PB, data collection: EL, data analysis: EL, DD, manuscript preparation: EL, PB, manuscript approval: all.

## Data availability

Table S1 with raw data

## SUPPLEMENTARY FILES

Table S1: table with phenotypic data for individual flies

Table S2: stats tables

Table S3: GWAS hits

Figure S1: thermal plasticity

Figure S2: Manhattan Plots

**Figure S1.**
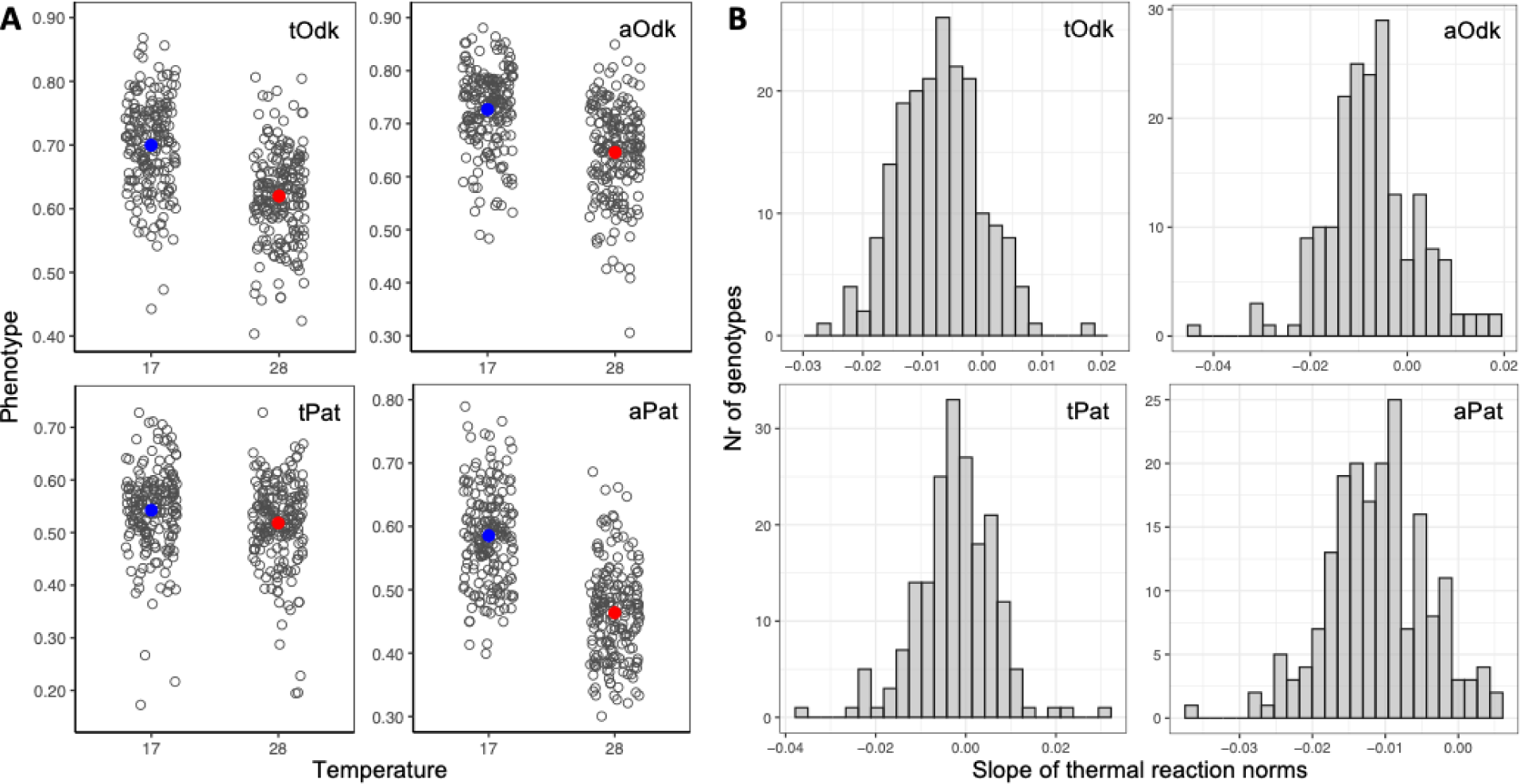
Thermal plasticity in body pigmentation. For each of the four target traits (tOdk, aOdk, tPat, aPat) the graphs display: **A)** phenotypes of all DGRP flies (one dot per fly) separated by developmental temperature (with mean values in color; blue for 17°C and red for 28°C developmental temperature); **B)** frequency distribution of slope of thermal reaction norms in the DGRP genotypes, illustrating bell-shaped distribution characteristic of quantitative traits.

**Figure S2.**
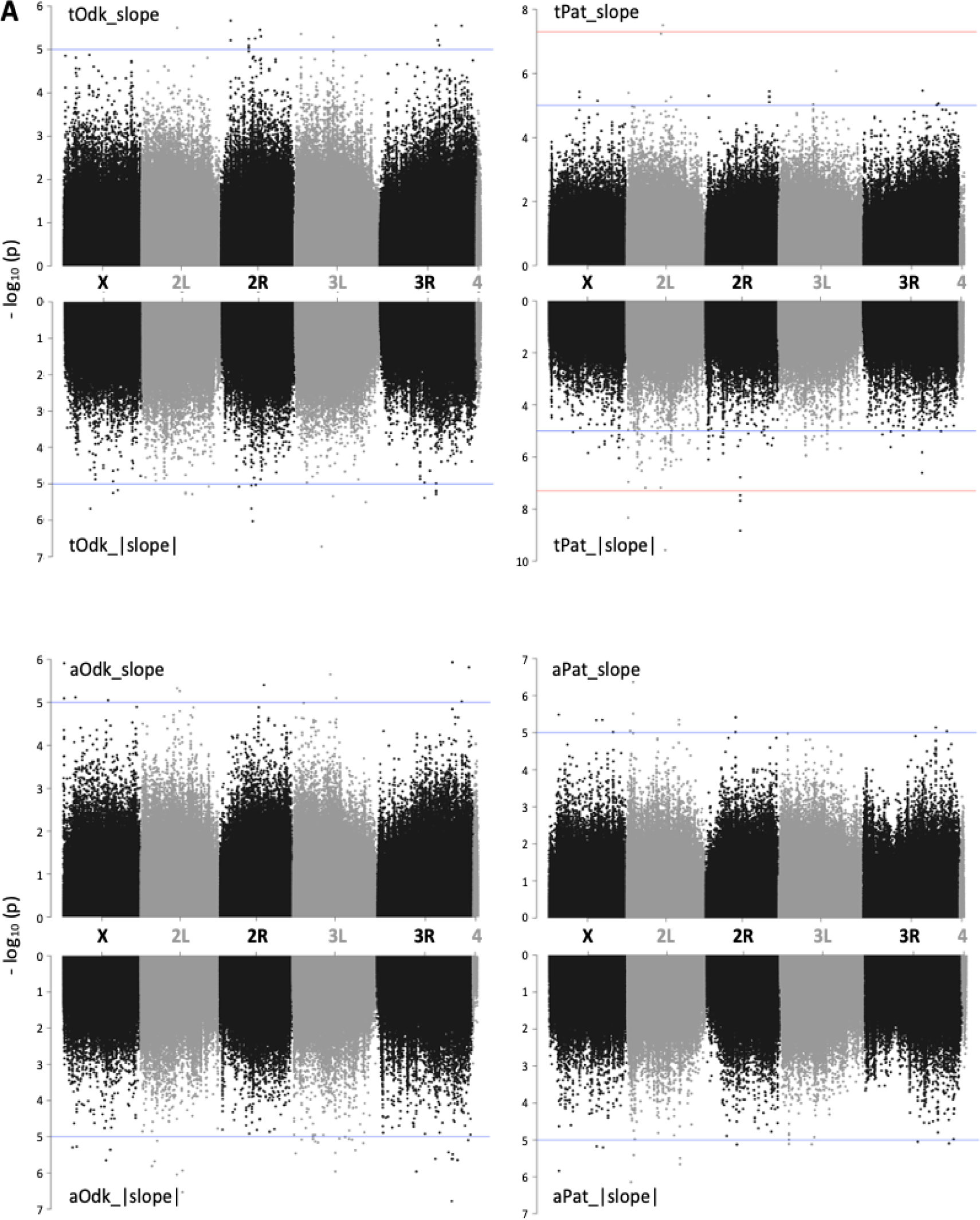
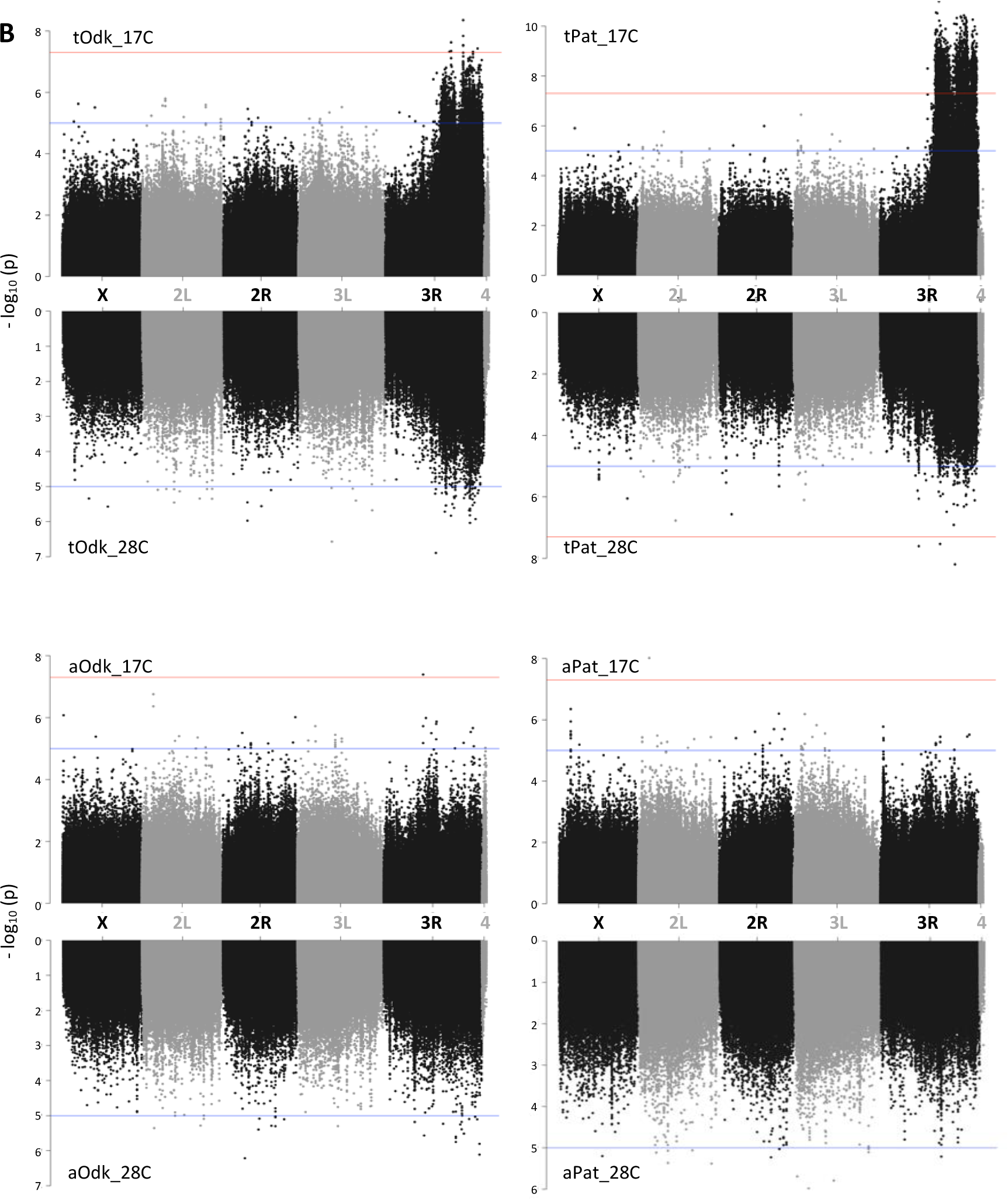
**Manhattan plots** corresponding to the GWAS for variation in different properties of our target pigmentation traits: tOdk and tPat (top), aOdk and aPat (bottom). **A)** variation in thermal reaction norms: slope (upwards) and |slope| (downwards). **B)** variation between individuals developed at each of the temperatures: 17°C (upwards) and 28°C (downwards). Horizontal lines represent threshold p-value of 10e-5 (blue) and 10e-7.5 (red).

