## Supplementary figures and images for "Genetic basis of variation in thermal developmental plasticity for *Drosophila melanogaster* body pigmentation"

### Figure S1

#### Figure S1. Thermal plasticity in body pigmentation

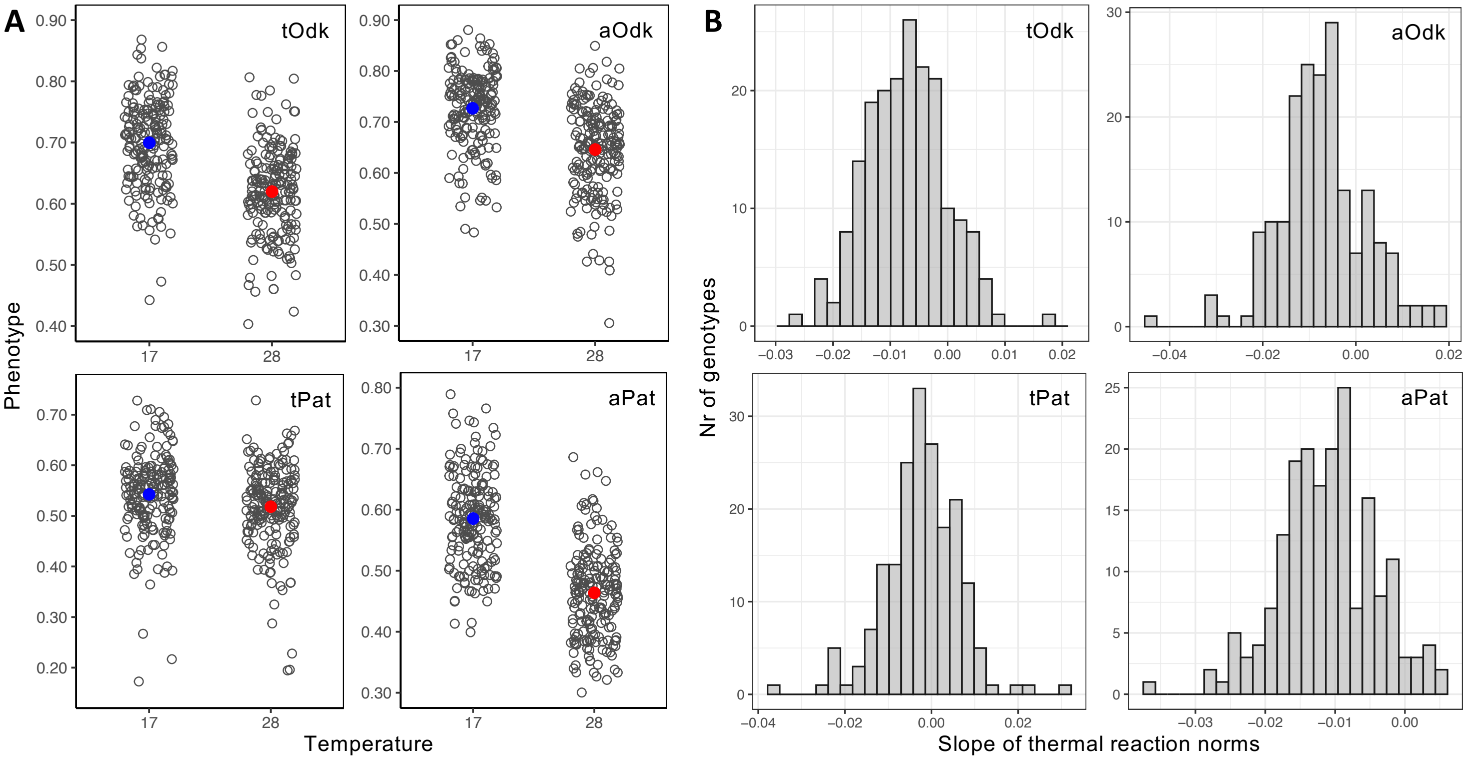

### Figure S2

#### Figure S2. MPs

A

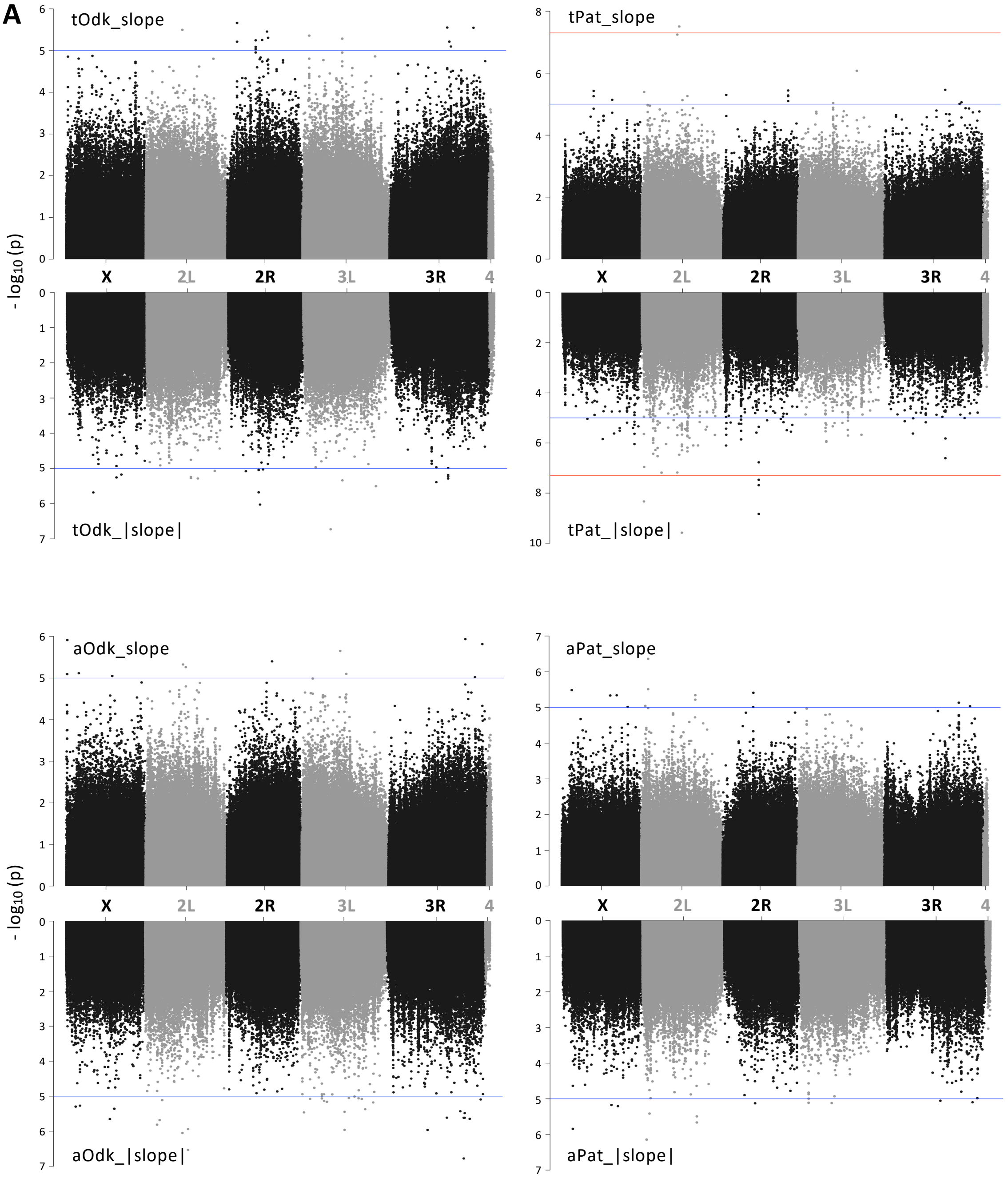

#### Figure S2. MPs

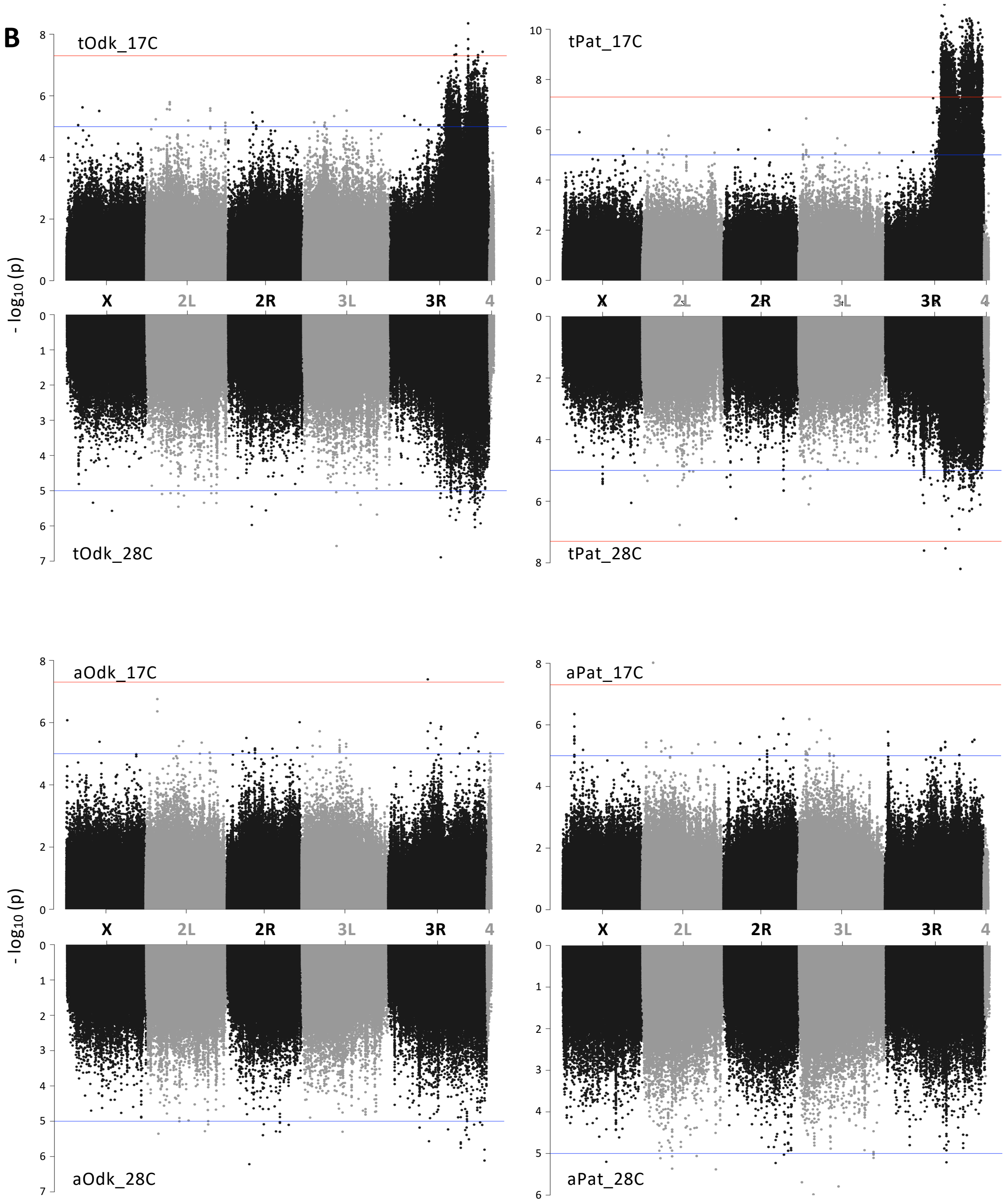
