## Supplementary material for "Genetic basis of variation in thermal developmental plasticity for *Drosophila melanogaster* body pigmentation": Table S2

**Pigmentation GWAS - Results from statistical analyses**

### **Linear models - GxE effects**

#### tOdk

*mod <- lm(Odk ~ DGRP_line * Temperature, data=allDGRPs_traits_T)*

*aov <- anova(mod)*

| **Response: Odk** | Df | Sum Sq | Mean Sq | F value |  | Pr(>F) |
| --- | --- | --- | --- | --- | --- | --- |
| DGRP_line | 199 | 17,4342 | 0,0876 | 48,642 | < | 2,20E-16 |
| Temperature | 1 | 5,9448 | 5,9448 | 3300,628 | < | 2,20E-16 |
| DGRP_line:Temperature | 190 | 6,0161 | 0,0317 | 17,58 | < | 2,20E-16 |
| Residuals | 4356 | 7,8456 | 0,0018 |  |  |  |

#### tPat

*mod <- lm(Pat ~ DGRP_line * Temperature, data=allDGRPs_traits_T)*

*aov <- anova(mod)*

| **Response: Pat** | Df | Sum Sq | Mean Sq | F value |  | Pr(>F) |
| --- | --- | --- | --- | --- | --- | --- |
| DGRP_line | 199 | 16,955 | 0,0852 | 8,3449 | < | 2,20E-16 |
| Temperature | 1 | 0,629 | 0,62896 | 61,6037 |  | 5,25E-15 |
| DGRP_line:Temperature | 190 | 8,655 | 0,04555 | 4,4616 | < | 2,20E-16 |
| Residuals | 4356 | 44,474 | 0,01021 |  |  |  |

#### aOdk

*mod <- lm(Odk ~ DGRP_line * Temperature, data=allDGRPs_traits_AP)*

*aov <- anova(mod)*

| **Response: Odk** | Df | Sum Sq | Mean Sq | F value |  | Pr(>F) |
| --- | --- | --- | --- | --- | --- | --- |
| DGRP_line | 199 | 19,9064 | 0,100 | 30,893 | < | 2,20E-16 |
| Temperature | 1 | 6,4116 | 6,4116 | 1980,084 | < | 2,20E-16 |
| DGRP_line:Temperature | 190 | 10,7995 | 0,0568 | 17,554 | < | 2,20E-16 |
| Residuals | 4521 | 14,6393 | 0,0032 |  |  |  |

#### aPat

*mod <- lm(Pat ~ DGRP_line * Temperature, data=allDGRPs_traits_AP)*

*aov <- anova(mod)*

| **Response: Pat** | Df | Sum Sq | Mean Sq | F value |  | Pr(>F) |
| --- | --- | --- | --- | --- | --- | --- |
| DGRP_line | 199 | 23,041 | 0,1158 | 32,0436 | < | 2,20E-16 |
| Temperature | 1 | 16,527 | 16,5268 | 4573,7514 | < | 2,20E-16 |
| DGRP_line:Temperature | 190 | 6,204 | 0,0327 | 9,0366 | < | 2,20E-16 |
| Residuals | 4521 | 16,336 | 0,0036 |  |  |  |
